# Is there a fly in my soup? To what extent do metabarcoding and individual barcoding tell the same story?

**DOI:** 10.1101/2025.11.12.687976

**Authors:** Brendan Furneaux, Tomas Roslin, Bess Hardwick, Deirdre Kerdraon, Hannu Autto, Gaia Banelyte, Jeremy R. deWaard, Stephanie L. deWaard, Arielle Farrell, Oula Kalttopää, Erik Kristensen, Hanna M. K. Rogers, Jayme E. Sones, Evgeny V. Zakharov, O. Ovaskainen

## Abstract

1. Metabarcoding has become the method of choice for characterising complex arthropod communities. The extent to which metabarcoded bulk samples will recover the same community composition as individual sequencing of all individuals in the sample remains poorly quantified. Biases such as unequal extraction of DNA from different taxa, primer mismatches and non-random PCR may cause the selective drop-out of species from metabarcoding data. At the same time, DNA metabarcoding may reveal arthropod taxa present not as individuals, but as DNA residues on the surface or in the gut of insects.
2. To quantify the consistency in sample contents established by different means, we metabarcoded 45 bulk insect samples, then extracted all arthropods and sequenced them individually. Metabarcoding targeted 418 bp at the 3’ end of the Folmer barcoding region, while individual barcodes captured the entire 658 bp Folmer region. The metabarcoding workflow, including PCR amplification, sequencing, and bioinformatics, was performed in three replicates from three separate lysate aliquots per sample. Sequences were assigned to Barcode Index Numbers (BINs) as identical taxonomic categories across data types.
3. We found that metabarcoding is replicable, as different replicates of the same sample recover similar species richness and composition. Individual barcoding and metabarcoding provide the same impression of relative differences in community structure: estimates of relative species richness and relative dissimilarity between sample pairs are congruent among data types. Dissimilarity between data types varies with BIN richness in the sample, but this relationship reflects nestedness rather than turnover: metabarcoding recovers the same set of core species as individual barcoding but adds hundreds of species on top. Any BIN recovered as an individual occurred with high probability in the metabarcoding data, and any BIN found in high read abundances by metabarcoding was likely found as an individual.
4. Our analysis suggests that metabarcoding data will closely mimic the sample contents in terms of arthropod species richness and composition. Taxa recovered in low copy numbers in metabarcoding sequence data will typically represent DNA left as residues from past biotic interactions. Barring sequencing errors, both types of data yield biologically relevant insights into the taxa present in the source community.

## Introduction

Metabarcoding of arthropod samples is rapidly becoming the go-to method for biodiversity discovery (Cristescu, 2014; Souto-Vilarós et al., 2024; Yu et al., 2012), community comparisons (Barsoum et al., 2019; Taberlet et al., 2012), ecological monitoring (Granqvist et al., 2025; van Klink et al., 2022, 2024) and testing of topical theories in community ecology (Abrego et al., 2021; Andújar et al., 2022). Indeed, metabarcoding is arguably the only current method allowing us to thoroughly characterise large and complex arthropod samples at a relevant scale, such as the ca. 20,000 Malaise trap samples collected by the LIFEPLAN project (Hardwick et al., 2024; Ovaskainen et al., 2025). Yet, all new types of data come with both benefits and caveats, and metabarcoding data are no exception (Hartig et al., 2023). A concern repeatedly raised is the extent to which metabarcoding will suffice to reveal *all* taxa actually present in the sample, i.e., the “true sample content” – typically taken to be the species present as recognizable individuals (Basset et al., 2022; Elbrecht et al., 2017; Yu et al., 2012). When metabarcoding fails to detect species which might, in principle, have been detectable by a (very large) set of skilled taxonomists or the (expensive and tedious) individual barcoding of all individuals present in the sample (as done in Global Malaise Trapping Project; Geiger et al., 2016; Seymour et al., 2024), then the full method is oftentimes discharged as being “incomplete” or “biased”. When samples reveal taxa not present as individuals, they are taken to represent “contamination”, “misidentifications” or “false positives” (Iwaszkiewicz-Eggebrecht et al., 2023; Ji et al., 2020).

There are many reasons why species present in the sample as individuals may (and typically will) be lacking from metabarcoding data (Lamb et al., 2019). Well-known and amply studied reasons include unequal extraction of DNA from taxa of different structure (Braukmann et al., 2019; Marquina et al., 2022), primer mismatches and PCR biases (Elbrecht et al., 2019a; Piñol et al., 2019), and the swamping of signal from rare species by more abundant ones in sequence yields of realistic size (Elbrecht & Leese, 2015). Each of these factors may result in the systematic underrepresentation of selected taxa, thereby causing a bias in the data and downstream analyses. The basic chemistry of PCR and sequencing reactions are also associated with stochasticity, generating random noise (Leray & Knowlton, 2017; Shelton et al., 2023). Given the large number of DNA templates present in a complex sample, some templates may get amplified by chance alone, whereas others fail to do so (Piñol et al., 2019). As a result, samples processed in replicates typically show some differences in the exact species pool recovered (Van Den Bulcke et al., 2021; Zinger et al., 2019), with individual species “dropping out” from the sequence yield of individual replicates.

The challenge of “false negatives” (i.e., species present in the sample but not detected in the metabarcoding data) has been studied *in silico* (Ficetola et al., 2015, 2016; Lahoz-Monfort et al., 2016), *in vitro* (Piñol et al., 2015), and empirically, through the creation of so-called “mock communities” of known contents (Elbrecht et al., 2019b; Elbrecht & Leese, 2015; Martoni et al., 2022; Yu et al., 2012). Such communities are typically assembled from a handful to some tens of species (Blanckenhorn et al., 2016; Iwaszkiewicz-Eggebrecht et al., 2023; Piñol et al., 2019) (but for more ambitious approaches, see Creedy et al., 2019; Elbrecht et al., 2019b). A common finding is that detection rates are never perfect. Some taxa (such as ants; Basset et al., 2022; Iwaszkiewicz-Eggebrecht et al., 2023) appear to be particularly hard to detect – and poor detectability can sometimes be attributed to simple correlates, including a hard and sclerotized surface (Iwaszkiewicz-Eggebrecht et al., 2023; Marquina et al., 2022).

Yet, there are also perfectly good reasons why species *not* present in the sample as individuals will appear in the sequence yield. In particular, a large number of studies have shown how individual arthropods will accumulate DNA traces of past interactions (Borsato et al., 2025; Wirta et al., 2022), from DNA in gut contents (Eitzinger et al., 2019; Kocher et al., 2017; H. Wirta et al., 2015) to DNA from past hosts or current parasitoids (Wirta et al., 2014). Critically scanning for “false positives” (i.e., species appearing in the sample even when no one put them there) will thus be an unrewarding task – unless the community was generated from laboratory-reared individuals fed on DNA-free food.

Given the challenges involved both in establishing biologically-relevant mock communities, and in defining the ground-truth based on *either* metabarcoding data *or* individually-sequenced specimens, we suggest a more constructive approach: to quantify the factors affecting the probability by which a taxon detectable as an individual in the sample is also detected in the metabarcoding data, and – *mutatis mutandis* – the probability with which a taxon detected in the metabarcoding data will also be present as an individual. In this paper, we draw on a set of 45 samples exhaustively characterized by both metabarcoding and sequencing of individuals to achieve this. More specifically, we ask:

1. Is metabarcoding in itself replicable: do different replicates of the same samples recover the same species richness and the same species composition from a sample?
2. Does individual barcoding and metabarcoding provide the same impression of community structure:
  2a. Do estimates of species richness concur among methods?
  2b. Do estimates of compositional differences between samples concur among methods?
  2c. Does the concurrence between data types depend on Barcode Index Number (BIN) richness in the sample?
3. Can we predict the probability of species occurrence in one type of data by the BIN’s abundance in the other type (*sensu* individuals for individual barcodes and read counts for metabarcoding data)?

## Materials and methods

Forty-five insect samples were collected by Malaise traps across 19 sites of which 18 were located in Sweden, and the last near the border in Finland. All data and the underlying methods were released by Orsholm et al. (2025). In brief, we first metabarcoded the samples following the protocol outlined by deWaard et al. (2024). To assess the reproducibility of metabarcoding, we split the DNA extract from each bulk sample into three replicates, and processed these independently through PCR, library preparation, sequencing, and bioinformatics. We then exhaustively DNA barcoded all detectable arthropods individually following Ivanova et al. (2006) for DNA extraction and Hebert et al. (2018) for sequencing. Barcode sequences from the individual specimens were added to BOLD (Ratnasingham & Hebert, 2007), where they were assigned to existing barcode identification numbers (BINs) or being used to delineate new BINs if they did not match any existing BIN. Taxa were delimited for the metabarcoding sequences using the BIN matching algorithm of mBRAVE (Ratnasingham, 2019), with the individual-level sequences produced in this study included in the reference database. We discarded sequences from the metabarcoding dataset which were not mapped to BINs assigned to phylum Arthropoda. Because BINs are at present only generated for sequences derived from individual specimens, this means that all sequences retained for analysis can in principle be linked to a physical specimen, whether from the present study or another. Thus, we adopted BINs as our operational species definition. As a result of these choices, any taxa appearing in the metabarcoding and individually sequenced data were defined by the same criteria and directly matchable between the two data types. At the same time, the species richness recorded in the metabarcoding is likely to be an underestimate, because we included only species that could be mapped to existing BINs rather than e.g. clustering the raw sequences.

To evaluate the consistency between replicates and sequencing methods, we used as metrics species richness, and pairwise dissimilarity in species composition by Sørensen’s and Simpson’s indices. Sørensen’s dissimilarity is defined by (*b* + *c*)/(2*a* + *b* + *c*), where *a* is the number of BINs detected in both replicates, and *b* and *c* are the numbers of BINs detected in only the first or only the second replicate, respectively (Sørensen, 1948). Thus, the dissimilarity is the number of BINs found in only one of the two samples, divided by the sum of the total number of BINs found in each sample. This index reaches a value of zero when all BINs are identical between samples and one when all BINs are different. Sørensen’s index is known to be sensitive to differences in BIN richness – since any species detected uniquely by one method (*b* or *c* in the formula above) tend to drive the value of the index towards its maximum value of 1. As the metabarcoding data tended to detect more species than individually sequenced samples, we wanted to assess whether metabarcoding resolved *the same plus more* species than those detected by individual barcoding, or *totally different* species than detected by individual barcoding. The first phenomenon is referred to as nestedness (Baselga, 2010, 2012), since it is best illustrated by the case where all species in the more species-poor sample are also included in the more species-rich sample (i.e., the species pools are perfectly nested).

The second case is referred to as turnover, since it focuses on differences (i.e., turnover) in species identity. To resolve between these scenarios, we drew on another measure of community composition, Simpson’s (1943) distance. This metric was originally designed to be insensitive to differences in species richness. It is defined as min(*b, c*)/(*a* + min(*b, c*)), with *a, b*, and *c* as defined above – i.e., the fraction of BINs in the replicate with fewer BINs which are unique to that replicate. Thus, it reaches a value of zero when the BINs in the less BIN-rich replicate are a subset of the BINs in the more BIN-rich replicate and a value of one when the two replicates have completely different BINs. In the case where the two replicates have the same total number of BINs, Sørensen’s and Simpson’s dissimilarities are equal (Baselga, 2010). Drawing on this rationale, we repeated the analyses described below using Simpson’s distance index. Sørensen and Simpson dissimilarities were calculated in R version 4.4.3.

To assess the replicability of metabarcoding in terms of species richness, we fitted a linear model of BIN richness explained by the sample (factor with 45 levels). The model was fitted in R using function lm, and we measured replicability by the R^2^ of this model. To assess the replicability of metabarcoding in terms of species composition, we compared the Sørensen and Simpson dissimilarities between different replicates deriving from each sample to the dissimilarity between these and replicates deriving from other samples.

To assess the comparability of metabarcoding and individual barcoding in terms of species richness, we fitted a linear model of BIN richness detected by metabarcoding as a function of the BIN richness recovered by individual barcoding. Here, the intercept (if positive) is an estimate of how many arthropod BINS were detected by metabarcoding in samples that had a very low (zero) number of BINs detected as individuals. The slope reveals how many new BINs we detect by metabarcoding for every new BIN detectable by sequencing of individuals. If each BIN were detected by both methods, then we would expect an intercept of zero and a slope of one. The R^2^ of the linear model indicates how well we may predict species richness detected by one method based on the species richness detected by the other method. The model was fitted in R using function lm. To assess the comparability of metabarcoding and individual barcoding in terms of species composition, we computed the Sørensen and Simpson dissimilarities between the individual-level results from a sample and the three metabarcoding replicates from the same sample. We then used a Mantel test to compare the observed Pearson’s product-moment correlation coefficient, *r*, between the observed, pairwise dissimilarities to the expected distribution of *r* when the data were reshuffled among sample pairs within the other data type. This analysis was implemented in R version 4.4.3 using package vegan (Oksanen et al., 2025) with 999 permutations. As we expected that low BIN richness would lead to larger nestedness, we tested whether the Sørensen and Simpson dissimilarities between the two sampled types covaried with the BIN richness resolved by barcoding of individuals or by metabarcoding. Since the two methods detected slightly different numbers of BINs, we evaluated patterns against both BIN richness resolved by individual barcoding and BIN richness resolved by metabarcoding.

Finally, we assessed how the abundance of a BIN in one type of data was reflected in its detection probability in the other type of data – e.g., whether taxa found to be rare in one type of data are likely to be missed by the other type of data. For each data type (metabarcoding or individual barcoding), we fitted a logistic regression model of the presence/absence of a BIN in data generated by one method as a function of the log(number of individuals or reads) by the other method. Each BIN in each sample was used as an individual data point. Here, the intercept will reflect the basic probability that a BIN present in a sample of one data type will be detected by the other. When metabarcoding data are used as the response, the slope will reflect how quickly this probability increases with logarithmically more individuals in the sample. When individual barcoding data are used as the response, the slope will reflect how quickly this probability increases with logarithmically more reads in the sequence yield. In predicting the presence of a BIN among metabarcoding reads from BIN presence among individual sequences, we added log(sequencing depth) as a covariate. To see why, consider a sample with low sequencing depth (i.e., few reads in total). For such a sample, all reads – regardless of whether the BIN is present as an (or several) individual(s) or not – will be rarer than in a sample of deep sequencing depth, and the probability of a BIN yielding zero reads will thus be higher. All models were fitted in R using the function glm.

## Results

The metabarcoding resulted in a total of 29.1 million reads, with an average of 214,000 reads per sample replicate (SD 59,000, range 89,000– 361,000). When resolved by metabarcoding, individual sample replicates contained between 84 and 697 BINs (mean 250 ± SD 144).

A total of 36,402 individual arthropods were sorted from the samples (per-sample mean 809 ± SD 750, range 36–3,228). Of these, 32,598 (89.6%) successfully yielded an individual barcode sequence which passed standard quality controls (per-sample mean 724 ± SD 651, range 32–2,538), including an order-level match to the morphological taxonomic designation provided during sorting. When resolved by individual sequences, samples contained between 10 and 749 BINs (mean 174 ± SD 172).

Metabarcoding turned out to be highly replicable, as species richness was highly consistent among replicates of the same sample (R^2^ = 0.997). Also species composition was replicable in the sense that it varied much more between samples than between replicates of the same sample. On average, Sørensen’s index of dissimilarity between replicates from the same sample was 0.23 ± 0.08 (see marginal distributions in Fig. 1A,B), and Simpson’s distances were similar, mean 0.21 ± 0.07 (marginal distributions in Fig 1C,D). In contrast, Sørensen and Simpson dissimilarities between replicates from different samples were substantially higher; 0.86 ± 0.06 and 0.80 ± 0.06, respectively (Fig. 1).

**Figure 1.**
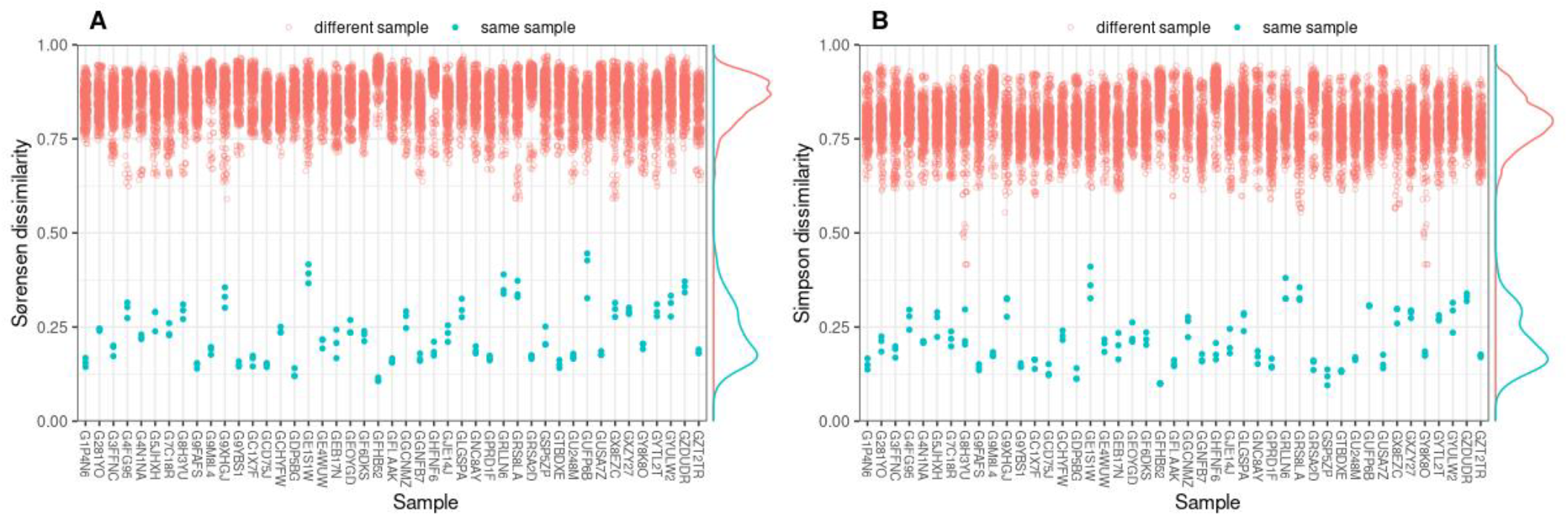
Replicability of metabarcoding samples. Replicability is based on samples processed in three replicates and measured through measured by Sørensen (A) and Simpson (B) dissimilarities. Blue points refer to comparisons between replicates of the same sample, red points to comparisons to other samples.

The species richness recovered was also highly consistent among methods, in the sense that a sample perceived to be rich in BINs by one method was also rich in BINs by the other (Fig. 2). Metabarcoding resolved many taxa also from samples for which we detected a low species richness by individual barcoding (intercept = 172±SE 10.9), but for each new BIN resolved by the sequencing of individual insects we detected less than one new BIN by metabarcoding (slope = 0.94±SE 0.04). Although the exact intercept and slope values varied, these patterns remained constant when considering each metabarcoding replicate separately (Fig. 2A) as when pooling replicates within samples (Fig. 2B), or when considering only BINs present in all three replicates of each sample (Fig. 2C).

**Figure 2.**
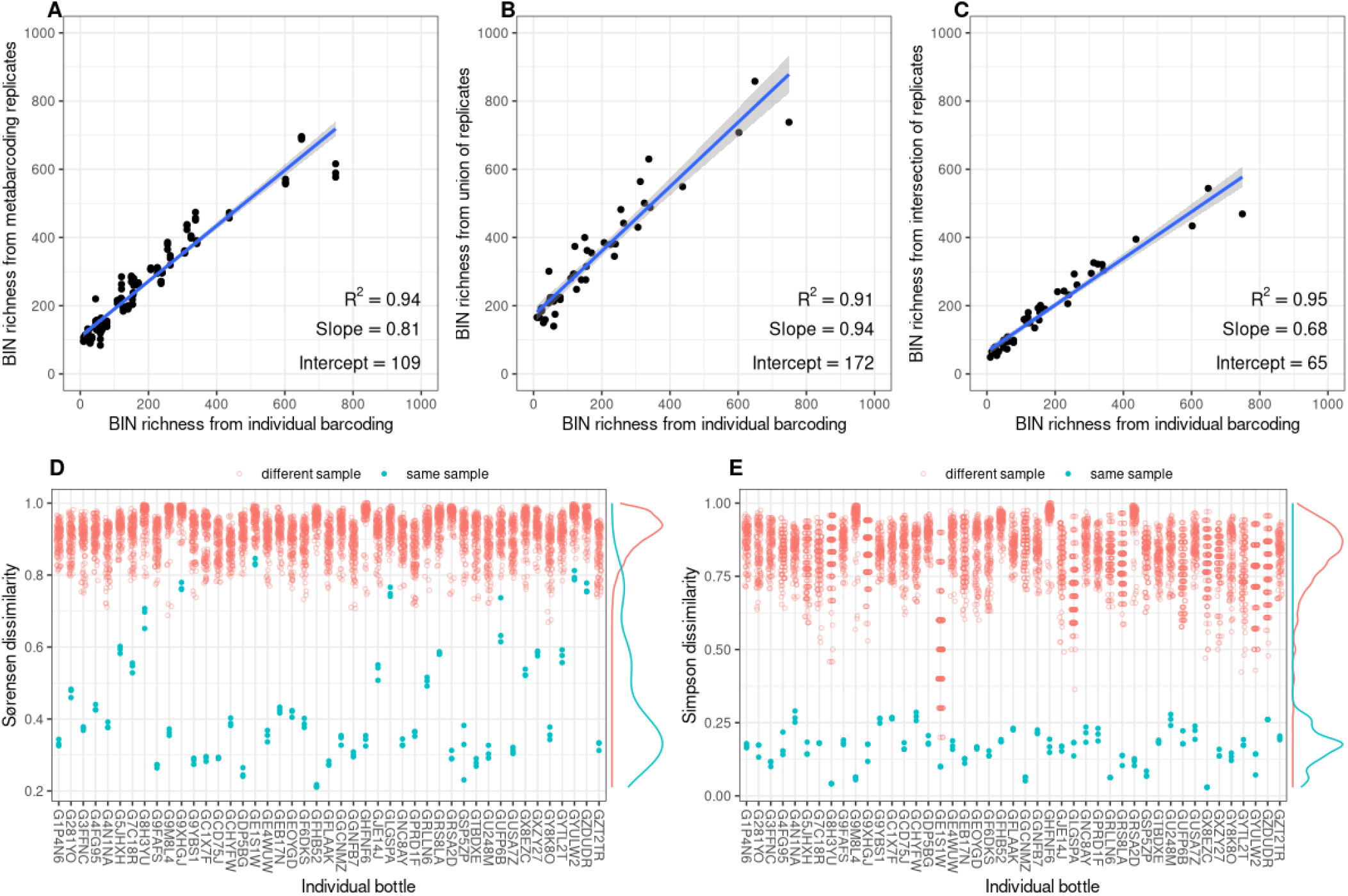
Comparison between individual barcoding vs. metabarcoding. The upper panels compare BIN richness revealed by individual barcoding (x-axis) to BIN richness revealed by metabarcoding (y-axis), either counting each replicate separately (A), pooling replicates within each sample (B), or including only BINs present in all three metabarcoding replicates of each sample (C). The line shows a linear model fitted to the data, of which key statistics are inserted in the panels. The lower panels show Sørensen (D) and Simpson (E) dissimilarities between individual barcoding vs. metabarcoding for rows of samples. For the individually sequenced data from each sample, we show dissimilarity with all metabarcoding replicates. Blue points refer to comparisons between observations of the same sample, red points to comparisons to other samples.

Not only species richness, but also the community composition recovered was highly consistent among individual barcoding and metabarcoding results for the same sample. This was especially the case with Simpson’s dissimilarity, for which dissimilarity between the two methods was on average 0.17 ± 0.06 within samples and 0.83 ± 0.10 between samples. Sørensen’s dissimilarity between the methods was on average 0.44 ± 0.17 within samples and 0.92 ± 0.05 between samples. The higher values of within-sample Sørensen dissimilarity (and hence, patterns of nestedness) were especially seen in samples with low BIN richness (especially as measured by individual sequencing, Fig. 3), whereas Simpson’s dissimilarity was independent of BIN richness. The consistency between the methods in their estimated community composition was also reflected in the observed correlation between dissimilarity among sample pairs: this was much higher both for Sørensen (*r*: 0.69; P=0.0001) and Simpson’s dissimilarity (*r*: 0.52; P=0.0001) than expected under random permutation.

**Figure 3.**
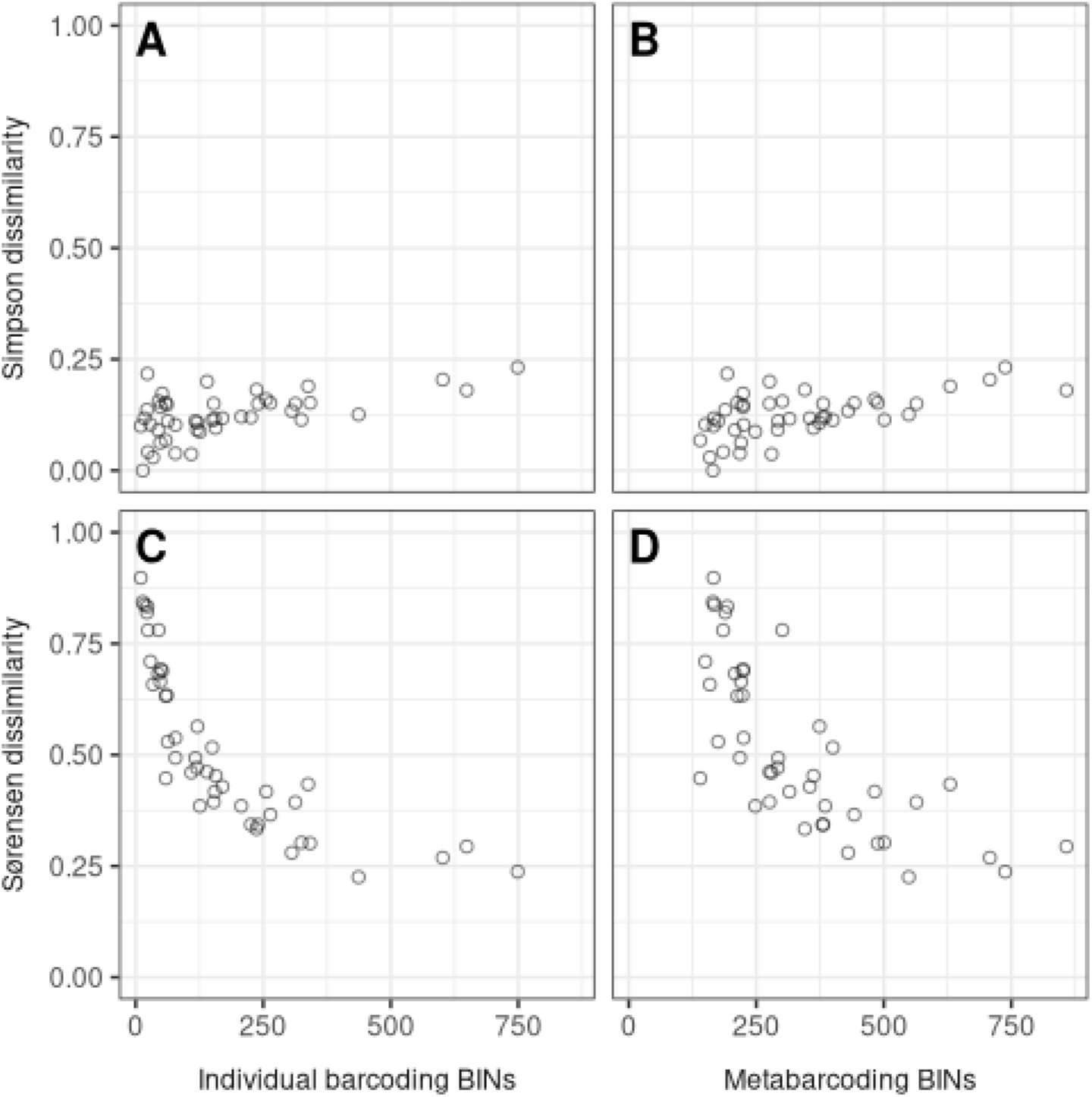
Dependency of between individual barcoding vs. metabarcoding dissimilarity on species richness. The panels show Sørensen dissimilarity (upper panels) and Simpson’s dissimilarity (lower panels) between community composition detected in individually sequenced and metabarcoded samples, plotted as a function of BIN richness in the sample. Since the two methods detected slightly different numbers of BINs (Fig. 2), we plot the same dissimilarity values against BIN richness resolved by either individual barcoding (left-hand panel) or metabarcoding (right-hand panel). Each data point is an individual sample, with the dissimilarity between the community detected by the two data types shown by the y-axis.

The probability of detecting a BIN by metabarcoding was high (70% for median metabarcoding sequencing depth) even for BINs found as a single individual during individual barcoding (Fig. 4). From this value, it further increased both with the abundance of the BIN in the individual barcoding data and with increased metabarcoding sequencing depth (Fig. 4A). Conversely, a BIN present as a single metabarcoding read was unlikely (13% probability) to be present as an individual (Fig. 4B). The probability of presence as an individual increased in a highly predictable manner with the number of metabarcoding reads, reaching 50% when the number of metabarcoding reads equalled 60 and reaching 95% when the number of metabarcoding reads equalled 50,000. At the level of individual arthropod orders, these patterns were qualitatively consistent with Fig. 4 but showed some idiosyncrasies between given taxa (for full patterns, see Appendix S1, Fig. S1.)

**Figure 4.**
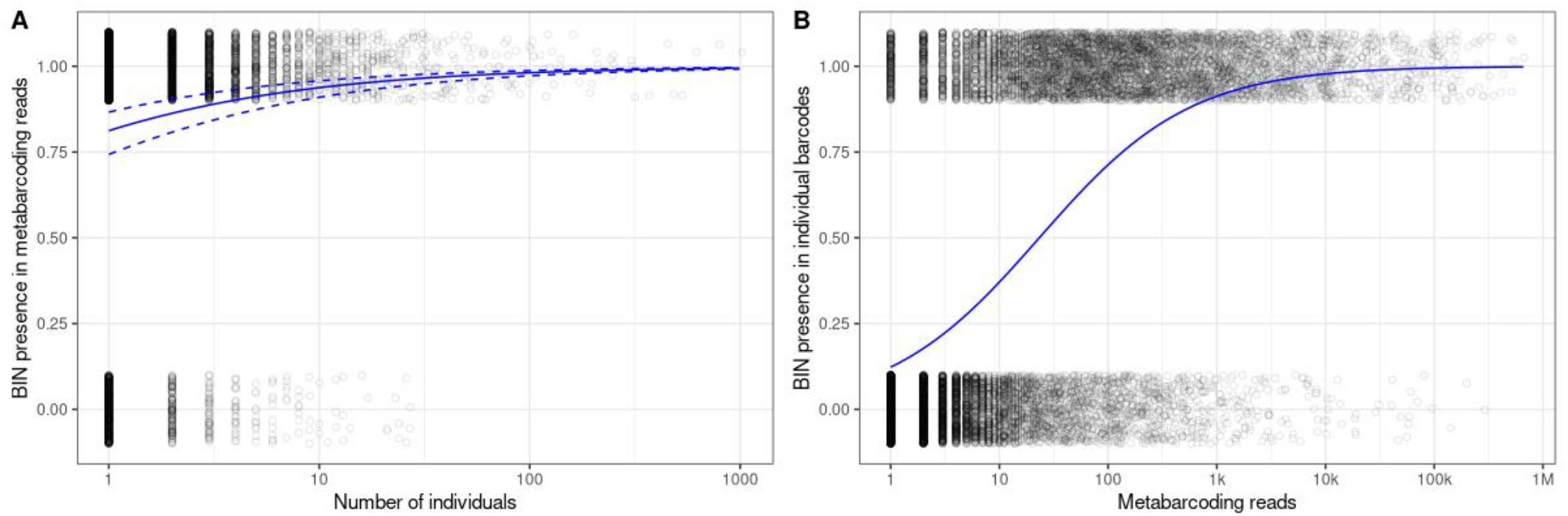
The probability of detecting a BIN present in one type of data in the other type of data. Panel (A) shows metabarcode presence as a function of individual abundance, and panel (B) individual presence as a function of the read abundance of this BIN in the metabarcoded data. The lines show a logistic regression model of Y regressed on X. In panel (A), the logistic regression model also includes log(metabarcoding sequencing depth) as a predictor, and the three lines show the prediction for sequencing depth of 500,000 (continuous line), 250,000 (lower dashed line), and 1,000,000 (upper dashed line) of this predictors. To resolve individual data points, presence/absence is plotted with jitter around 0 and 1.

## Discussion

While metabarcoding is becoming a method of choice for the characterisation of arthropod communities, its reliability remains a topic of controversy. In this paper, we have argued that neither sequencing of individuals nor metabarcoding will offer an objective ground truth for the DNA contents of an arthropod sample. To offer a nuanced approach, we asked a series of questions regarding the consistency of metabarcoding replicates, and the consistency of community structure gleaned from individual barcoding and metabarcoding data. We find that metabarcoding in itself is replicable, as different replicates recover similar species richness and composition from the same sample. When compared to individual barcoding, metabarcoding provides the same impression of relative differences in community structure, as estimates of species richness are highly correlated among methods, and as relational dissimilarity between samples prevail among data types. Importantly, the differences in species composition perceived between sample types do depend on BIN richness in the sample, but this relationship seems due to nestedness rather than turnover. Our most important findings relate to the predictability of BIN occurrence in one sample type by the other: here, any BIN recovered from individually sequenced arthropods was detected with high probability in the metabarcoding data, and any BIN found in high read abundances in the metabarcoding data was likely to be found as an individual. Below, we will discuss each finding in turn.

### Metabarcoding is qualitatively replicable

Different replicates of the same samples recovered much of the same species composition. An average of 77% of the BINs detected in any one replicate was shared with any other replicate from the same sample. When combined with the high probability (80%) that any BIN present as an individual in the sample was also detected in the metabarcoding sequence yield, these results attest to the replicability of metabarcoding assays and the robustness of the descriptors attained. Having said that, we fully subscribe to the general recommendation that metabarcoding should be run as replicates (Zinger et al., 2019), providing not only more data but also enabling quality control. Since chance variation will generate differences between replicates, more is probably better (Leray & Knowlton, 2017; Shelton et al., 2023). However, a counterpoint is offered by Smith & Peay (2014), who suggest that many studies which recommend PCR replicates are confounding replication with total sequencing depth. This confounding occurs as recommendations of replication are based on simply finding more of the low-read-abundance species from a mock community by sequencing 10 replicates, whereas no test is offered for whether the same study would have produced the same result by sequencing a single replicate 10 times deeper. The latter approach is arguably cheaper and faster, but allows fewer opportunities for quality control. It is also worth noting that the sequencing depth of individual replicates in this study (mean 214,000 reads) is substantially larger than used in many earlier studies, including Smith & Peay (pools of 1-16 replicates rarefied to 500 reads for 454, 38,000 reads for Illumina) or Leray & Knowlton (mean 65,000 reads per replicate), and this may be an important factor for replicability.

When samples are sequenced in replicates, the question emerges whether the data from individual replicates should be retained as nested data points of the same sample or pooled for analysis (as we have done in our subsequent analyses). This will depend on the modelling approach chosen, whereas for the current argument, the key finding is that individual replicates will be largely consistent, and the core species pool will be shared between individual replicates. That such consistency is the rule is also shown by another consideration: as a part of quality control, REF tends to omit taxa found in less than two out of three replicates, but this will typically result in the removal of less than 15% of BINs.

Overall, our results are consistent with those of Van den Bulcke et al. (2023), who showed high consistency in species numbers, Shannon indices and Inverse Simpson indices recovered for samples of marine benthos – even when the samples were analysed by different laboratories in a ring test.

### Individual barcoding and metabarcoding provide the same impression of relative differences in community structure

Our comparison of community metrics between metabarcoded and individually barcoded samples revealed a general agreement in terms of relative differences between the samples: a sample comparatively rich in BINs scored by one method was also rich in BINs scored by the other, and a sample dissimilar from other samples within one data type was also dissimilar within the other data type. This attests to the reliability of both methods, and to their partial interchangeability with each other.

In showing that metabarcoding can reliably recapture the contents of insect individuals visible in an arthropod sample, our findings support those of Remmel et al. (2024). These authors compared the arthropod contents of 12 Malaise samples as recovered by metabarcoding to the contents scored by morphological identification by taxonomic experts. Of 252 species identified in total, 54.8% were identified by both methods, whereas only 21.4% and 19.8% were detected exclusively by metabarcoding or morphology, respectively. Matching our current results, these authors also found good correspondence between methods in terms of total taxonomic richness per sample. In terms of community composition, our results echo the original findings of Yu et al. (2012). Using early 454 pyrosequencing techniques, these authors demonstrated that metabarcoding allows for precise estimation of both alpha diversity and pairwise community dissimilarity (beta diversity). Together, our studies thus show that metabarcoding will provide an accurate overview of the composition of local insect communities and differences between them, and that the patterns resolved are comparable to those determined through traditional, morphological identification or through individual barcoding.

Nonetheless, two differences do stand out between the metabarcoding and individual sequencing data here examined: First, the metabarcoded samples showed substantially higher species richness, with an average excess of 172 BINs for any one BIN detected by sequencing of individual insects (Fig. 2b). Second, starting from this initial excess (i.e., the intercept+1 in Fig. 2b), metabarcoding data accumulated 0.94 new BINs for each new BIN detected in individually sequenced samples (i.e., the slope in Fig. 2b).

Importantly, the higher BIN richness observed in metabarcoding data refers to arthropod BINs alone, since all other phyla were pruned from the data (see Methods). Thus, the higher BIN richness observed in metabarcoding data cannot be attributed to the resolution of a wide array of protists, bacterial symbionts etc. from the arthropods by metabarcoding data (Łukasik & Kolasa, 2024). Instead, a metabarcoded sample will reveal the presence of hundreds of *arthropod* species beyond the individuals trapped in the collecting bottle. This added set of species recorded seems attributable to a strong imprint of biotic interactions on the DNA content of an arthropod sample. Beyond the arthropods recovered from individually sequenced samples, each individual seems to carry a wide array of DNA residues from past encounters with other arthropods. We shall return to this issue below, in the context of predicting the probability of species occurrence in one type of data by BIN abundance in the other type of data.

When it comes to the slope of less than one (0.94) new BIN added to metabarcoding data for every new BIN detected by the sequencing of individuals, we note that this estimate is qualitatively consistent with a simple consideration: while arthropod communities are diverse, they are not limitless in their composition. Thus, towards more species-rich samples, the increase in the Sørensen index decelerates (Fig. 3). This may be due to much of the additional diversity (in the form of gut contents and surface residues) already being recorded from smaller samples, and richer samples thus showing higher redundancy with less species-rich samples (i.e., nestedness) – as there are simply less species left to add.

In terms of what BINs are resolved by the two data types, the difference in patterns between Sørensen and Simpson dissimilarities sheds light on the underlying phenomena (Fig. 3): while Sørensen dissimilarity increases among data types with increasing BIN richness in the target sample, no such relation was detected for Simpson dissimilarities (cf. Fig. 3A,B vs C,D). Since Sørensen dissimilarity is sensitive to differences in species richness but Simpson dissimilarities are not, the pattern in the former seems largely attributable to nestedness, i.e., to differences in BIN richness resolved by the two methods. Thus, metabarcoding seems to add further species beyond the core group occurring as intact individuals in the sample, whereas any BIN occurring as an intact individual is detected with a high probability (Fig. 4A).

### We can predict the probability of species occurrence in one type of data by its abundance in the other type of data

The probability of detecting a BIN present in one type of data in the other type of data, too, varied substantially with its abundance. We note that this effect is completely different from an effect of sequencing depth alone, since sample-to-sample variation in read depth was already captured by entering log(sequencing depth) as a covariate. Both before and after adjusting for this obvious effect, BINs more abundant in one type of data were more frequently detected in the other (Fig. 4). Nonetheless, the relationship differed substantially between when we used abundance among individuals sequenced to predict presence in metabarcoding data (Fig. 4A) and when we used metabarcoding read abundance to predict presence among sequenced individuals (Fig. 4B): even BINs present as single individuals in the sample were detected in the metabarcoding data with a probability of 83%, and this probability further approached one for abundant taxa (Fig. 4A). In contrast, BINs detected in low metabarcoding read numbers were unlikely to be found as recognizable individuals (P≅12%; Fig. 4A), whereas this probability quickly rose above 80% for BINs encountered in more than 1000 reads, and approached one for BINs encountered in more than 10,000 reads (Fig. 4A).

Overall, these findings show that any BIN collected as an entire individual in the bottle will be detected in metabarcoding data with a (very) high probability. Taxa rare in the sample will naturally be slightly harder to detect, as also found by Remmel et al. (2024). At the same time, our findings again attest to an excessive number of BINs *not* occurring as full individuals in the data, and thus *not* detectable by individual barcoding

– unless each individual is deeply sequenced and thereby essentially metabarcoded. The different relation uncovered when we used abundance among individuals sequenced to predict presence in metabarcoding data reveals a strong signature of biotic interactions on metabarcoded samples. As all the additional BINs will be arthropods, but arthropods not detectable as individuals, a major part is likely to emanate from gut contents. Here, both primary predation and even “secondary predation” (where a predator is consumed by another one after consuming a prey species; Tercel et al., 2021) seem likely to contribute. Malaise traps coupled with high throughput sequencing will thus reveal an even wider view of the local fauna than its content of insect individuals. Importantly, a species present in the gut content or on the surface of another will seem no less a member of the local community than a species present as an individual – and our current estimates suggest that the number of such species per physically detectable species ranges in the hundreds. Whether the sequencing of individual insects or metabarcoding will then constitute the preferred method (Chua et al., 2023) will depend on the question asked, but clearly, the two should be seen as mutually complementary rather than conflicting methods.

## Conclusions

Overall, our results provide a hope-inspiring view on metabarcoding as a method for generating reliable, verifiable and efficient insights into arthropod communities. Overall, the vast majority of taxa present will be detected by metabarcoding – with a probability greater than 80% even when a single individual is present. As such, it suggests that metabarcoding can actually deliver on the grand promise given by Ji et al. (2013) over a decade ago. At the same time, it is clear that metabarcoding involves some level of stochastic variation. Based on our results, the level of this stochastic variation is relatively small, and its effect can be mitigated by replication of the metabarcoding process. In our view, the most important take-home message from our study is that metabarcoding will resolve community features *beyond* those captured by individual barcoding, and that the two approaches should be seen as complementary rather than alternative to each other.

## Supplement S1. The probability of detecting a BIN present in one type of data in the other data type – split by arthropod orders

At the level of individual arthropod orders, relations between BIN occurrence in one data type and BIN abundance in the other data type were qualitatively consistent with overall patterns across taxa (see Fig 4 in the main text), but showed some idiosyncrasies between given taxa (Fig. S1). As examples, all Lepidoptera present in the sample were always likely to be detected in the metabarcoding data, whereas small mites were hard to detect (Fig. S1A). On the other hand, Opiliones (“daddy longlegs”) present in the metabarcoded data were rarely found as recognizable individuals (Fig. S1B).

That small mites are hard to detect is only to be expected, given that they include tiny amounts of DNA and can simultaneously be enclosed in sclerotized shells (Elo, 2019). Opiliones, on the other hand, feature extremely long and slender appendages, which are prone to breaking. Whether the frequent occurrence of daddy longlegs in metabarcoded data (Fig. S1B) will reflect that they often climb into the collecting part of the trap, but then escape after losing some appendage, and/or whether Opiliones are frequently consumed by predatory species later recovered in the traps cannot be established here – but both seem likely.

**Fig. S1.**
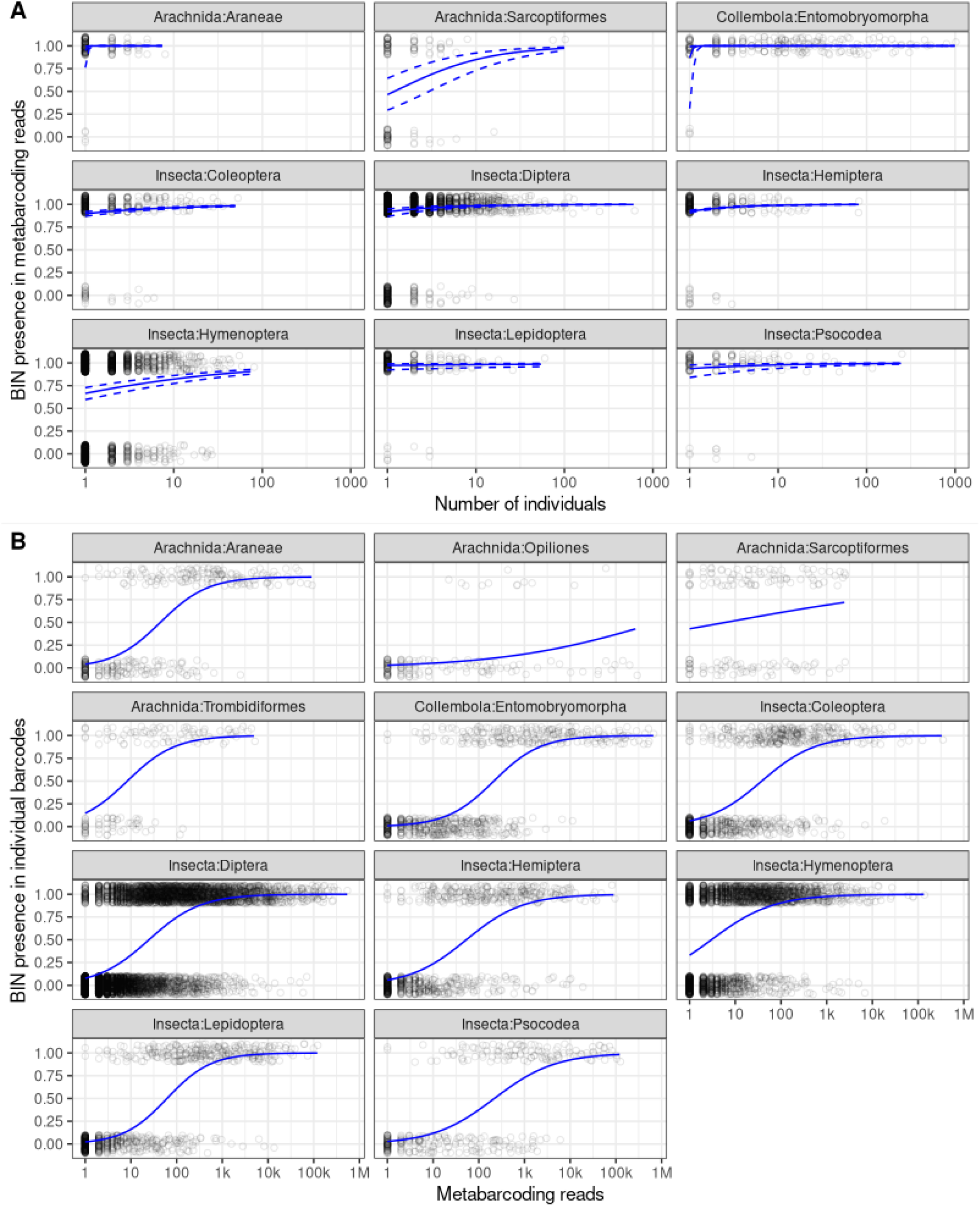
The probability of detecting a BIN present in one type of data in the other type of data modelled three ways: A) metabarcode presence as a function of individual abundance, B) individual presence as a function of the read abundance of this BIN in the metabarcoded data. Here, we break down the general pattern shown in Fig 4 of the Main text by arthropod orders. The lines show order-specific logistic regression models of Y regressed on X. To resolve individual data points, presence/absence is plotted with jitter around 0 and 1.

